# Double-strand break end configuration and 3D genome architecture are crucial for chromosomal translocation

**DOI:** 10.64898/2026.06.09.731072

**Authors:** Peifeng Liu, Jia Shou, Qiang Wu

**Author notes:** These authors contributed equally to this work (Peifeng Liu, Jia Shou).

## Abstract

Chromosomal translocations originate from erroneous rejoining of DNA double-strand break (DSB) ends and mechanisms of their formation remain largely unknown. Here we reveal DSB end configuration and three-dimensional (3D) genome architecture cooperatively shape both translocation frequency and junctional indel pattern. By integrating multiplex CRISPR/Cas9 editing with an optimized translocation assay, we find that translocation frequency correlates with spatial proximity. In addition, templated insertions emerge as a defining structural feature of translocation junctions induced by Cas9-mediated staggered cleavage and confirmed by DISCOVER-seq. Interestingly, translocation frequency is correlated with cleavage geometry. Finally, engineered Cas9 variants with altered DSB end configurations enable programmable modulation of junctional indel patterns. These findings shed significant insights into mechanisms of recurrent chromosomal rearrangements in human diseases and have interesting implications in safe applications of genome editing.

## Introduction

Chromosomal translocations arise from aberrant rejoining of DNA double-strand break (DSB) ends between non-homologous chromosomes and constitute a major class of genomic structural variation underlying cancers and inherited disorders (Gostissa et al. 2011; Lieber 2016). Such rearrangements perturb gene regulation and cellular identity by generating oncogenic fusion proteins or by rewiring enhancer-promoter interactions (Akdemir et al. 2020). Notably, cancer-associated translocations exhibit striking non-randomness, as exemplified by recurrent gene fusions of *BCR-ABL1, PML-RARA*, and *EWS-FLI1* in chronic myeloid leukemia, acute promyelocytic leukemia, and Ewing sarcoma, respectively (Daley and Baltimore 1988; Dethe et al. 1991; Delattre et al. 1992). The recurrent chromosomal rearrangements point to underlying molecular constraints, the nature of which remains insufficiently defined.

CRISPR/Cas9-mediated DNA-fragment editing with dual sgRNAs has emerged as a powerful strategy for engineering targeted chromosomal rearrangements, including deletions, inversions, duplications, and translocations (Jinek et al. 2012; Guo et al. 2015). Specifically, by simultaneously inducing DSBs at two genomic loci with dual sgRNAs, Cas9 harnesses endogenous cellular DNA repair pathways to religate distal DNA ends in non-native configurations, resulting in chromosomal rearrangements (Canver et al. 2014; Choi and Meyerson 2014; Maddalo et al. 2014; Guo et al. 2015; Kraft et al. 2015; Li et al. 2015; Shou et al. 2018; Wu and Shou 2020; Dahl-Jessen et al. 2025). In particular, when a pair of DSBs is generated on the same DNA molecule, repair can lead to excision of the intervening fragment or its reintegration but in the opposite orientation, generating deletions or inversions, respectively (Guo et al. 2015). By contrast, simultaneous Cas9 cleavages on different chromosomes can generate balanced or unbalanced translocations through interchromosomal end joining, enabling experimental modeling of oncogenic rearrangements such as the Philadelphia chromosome (Daley and Baltimore 1988). Initial studies suggested that the frequencies of chromosomal rearrangements are inversely related to the sizes of the intervening fragments (Canver et al. 2014). Subsequent studies, however, did not find the inverse correlation between rearrangement frequencies and the sizes of DNA fragments (Li et al. 2015; Binda et al. 2020). Importantly, the rearrangement outcomes of Cas9 editing are strongly influenced by cleavage geometry, local chromatin architecture, spatial proximity of breakpoints, and DNA repair pathway choice (Owens et al. 2019; Wu and Shou 2020; Kosicki et al. 2022). However, the exact mechanisms underlying chromosomal rearrangements are poorly understood.

In this study, we examined how 3D genome organization and DSB end configuration influence Cas9-induced chromosomal rearrangements. Using multiplex CRISPR editing in combination with an optimized chromosomal translocation assay (Hu et al. 2016; Yin et al. 2019; Liu et al. 2021), we mapped rearrangement junctions at single-nucleotide resolutions and found frequent templated insertions. Integration with high-resolution Micro-C maps reveals that translocation frequencies correlate with spatial contact frequencies. In addition, we showed that Cas9-generated staggered ends enhance translocation. Moreover, DISCOVER-seq via strand-resolved MRE11 ChIP-seq reveals asymmetric Cas9 cleavage *in vivo*. Finally, we demonstrated that engineered Cas9 variants enable programmable control of DSB end configuration and junctional indel pattern. Together, these findings suggested a unified framework in which 3D genome architecture and DSB end configuration jointly influence both the frequencies and junctional patterns of chromosomal translocations.

## Results

### Templated insertions as a junctional signature of Cas9-mediated chromosomal translocations

To investigate the molecular determinants of chromosomal translocation formation at single-nucleotide resolution, we initially employed a dual-sgRNA system (Guo et al. 2015; Li et al. 2015) to simultaneously induce DSBs at two different chromosomes and used an optimized PCR approach with a pair of primers to detect junctions of chromosomal translocations (Supplemental Fig. S1). Among the 90 translocations designed and assayed, we confirmed 77 correct translocation junctions by PCR cloning and Sanger sequencing (Fig. 1A; Supplemental Table S1). We then designed dual cleavages at clinically relevant breakpoints within the *HOXA6* (homeobox A6) and *BCR* (breakpoint cluster region) loci of chromosomes 7 and 22, respectively (Fig. 1B). Using an optimized Primer Extension-Mediated Sequencing (PEM-seq) approach (Hu et al. 2016; Yin et al. 2019; Liu et al. 2021), we captured all of the four rearrangement products: two derivative chromosomes corresponding to balanced translocations (der(7) and der(22)), a dicentric chromosome, and an acentric fragment (Fig. 1B,C). Synchronized induction of dual DSBs markedly increases translocation frequency compared with single DSBs (Fig. 1D; Supplemental Fig. S2A-C). Deep sequencing of all of the four translocation junctions revealed two prominent features (Fig. 1E-L; Supplemental Fig. S2D-K). First, all of the four translocation junctions exhibited extensive deletions flanking the breakpoints, whereas insertions were comparatively short (Fig. 1G,H; Supplemental Fig. S2F,G). Second, a subset of junctions contained non-random templated insertions precisely matching sequences adjacent to the cleavage sites (Fig. 1I-L). Specifically, der(7) junctions showed a strong bias toward T nucleotide insertions derived from the *BCR* site (Fig. 1I,K), whereas der(22) junctions preferentially incorporated “G” or “GAG” nucleotide insertions corresponding to the *HOXA6* flanking site (Fig. 1J,L). Dicentric junctions lacked a clear nucleotide bias (Supplemental Fig. S2H,J), while acentric junctions mirrored the T preference observed in der(7) (Supplemental Fig. S2I,K). These site-specific templated insertions (TINs) suggest that staggered cleavage by Cas9 facilitates templated nucleotide incorporation during the end-joining of chromosomal translocations.

**Fig 1.**
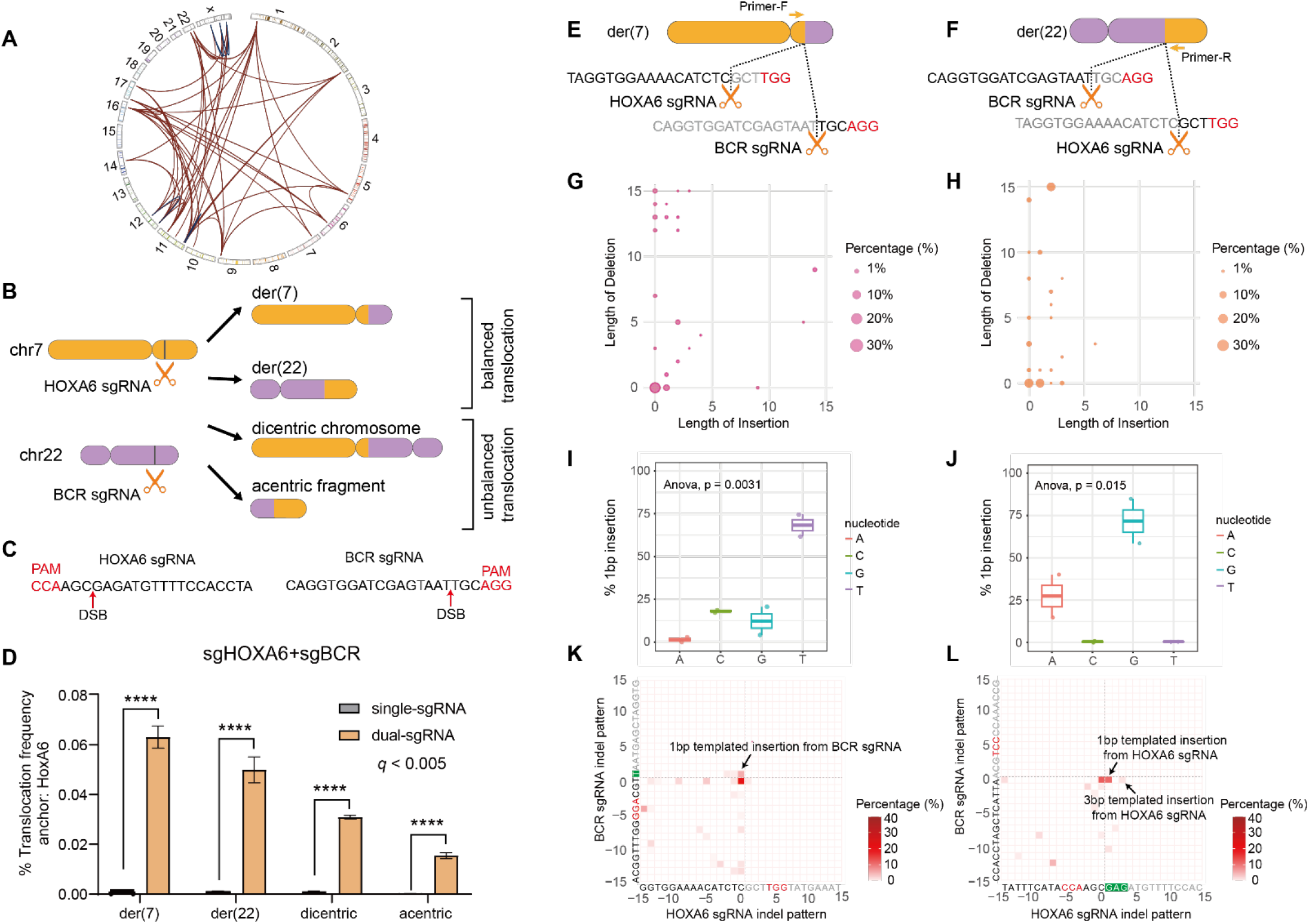
Templated insertions are characteristic junctional features at Cas9-induced chromosomal translocations. (**A**) Circos plot showing PCR assays of chromosomal translocations induced by Cas9 programmed with dual sgRNAs. (B) Schematic of the dual-sgRNA CRISPR/Cas9 system for chromosomal translocation. Simultaneous induction of DSBs at *HOXA6* (chr7) and *BCR* (chr22) by Cas9 programmed with dual sgRNAs generates four rearrangement products: der(7), der(22), a dicentric chromosome, and an acentric fragment. (C) SgRNA targeting sequences for *HOXA6* and *BCR* with PAM configurations and cleavage positions indicated by red arrows. (D) Frequencies of chromosomal translocations induced by single versus dual sgRNAs, normalized to the total number of detected events. (E, F) Schematic of balanced translocations (der(7) and der(22)), showing Cas9 cleavage sites, junction configurations, and detecting primer locations. (G, H) Bubble chart showing the frequency and length distributions of deletions and insertions at der(7) and der(22) junctions. Circle area is proportional to indel frequency; x-axis, insertion length; y-axis, deletion length. (I, J) Nucleotide composition of 1 bp insertions at der(7) and der(22) junctions. Box plots show median and interquartile range (Q1-Q3). (K, L) Heatmaps depicting the length and frequency distributions of deletions and templated insertions (TINs) mapped to the *BCR* and *HOXA6* cleavage sites. Zero denotes the sgRNA cleavage position; positive values indicate insertion length; negative values indicate deletion length; heatmap color intensity reflects indel frequency. Data in C are mean ± SD. Statistical significance was assessed using two-tailed Student’s *t*-test with Benjamini-Hochberg correction (D) and two-tailed one-way ANOVA (I, J). PAM sequences are highlighted in red. BCR, breakpoint clustered region; CRISPR, clustered regularly interspaced short palindromic repeats; Cas9, CRISPR associated protein 9; der(7), derivative chromosome 7; der(22), derivative chromosome 22; DSB, double-strand break; PAM, protospacer adjacent motif; sgRNA, single guide RNA.

### 3D genome architecture correlates with chromosomal translocation

To assess the role of nuclear organization in translocation formation, we induced DSBs at canonical *BCR-ABL1* breakpoints (*BCR* intron 14 and *ABL1* intron 1) to model the Philadelphia chromosome with *HOXA6* as an internal control (Fig. 2A-C; Supplemental Fig. S3A-D). We then simultaneously introduced breaks at *BCR, ABL1*, and *HOXA6* and quantified all of the four translocations between the *BCR* and *ABL1* loci with those between *BCR* and *HOXA6* as internal controls (Fig. 2D-G). Interestingly, we observed a significant increase of each of the four translocations between the *BCR* and *ABL1* loci compared to those between *BCR* and *HOXA6* (Fig. 2D,E). In addition, we observed a significant increase of each of the four translocations between the *ABL1* and *HOXA6* loci in comparison with those between *BCR* and *HOXA6* (Fig. 2F,G). Together, these observations suggest that there might be a translocation preference between *BCR* and *ABL1* loci. Indeed, integrating translocation frequencies with high-resolution Micro-C maps (Shanel et al. 2025) revealed that *BCR* and *ABL1* exhibit the closest spatial contacts among the three loci (Fig. 2H,I). Finally, quantitative analyses revealed a strong correlation between translocation frequency and spatial proximity (Fig. 2J; Supplemental Fig. S3E). All in all, these data suggest that 3D genome architecture plays an important role in chromosomal translocation.

**Fig 2.**
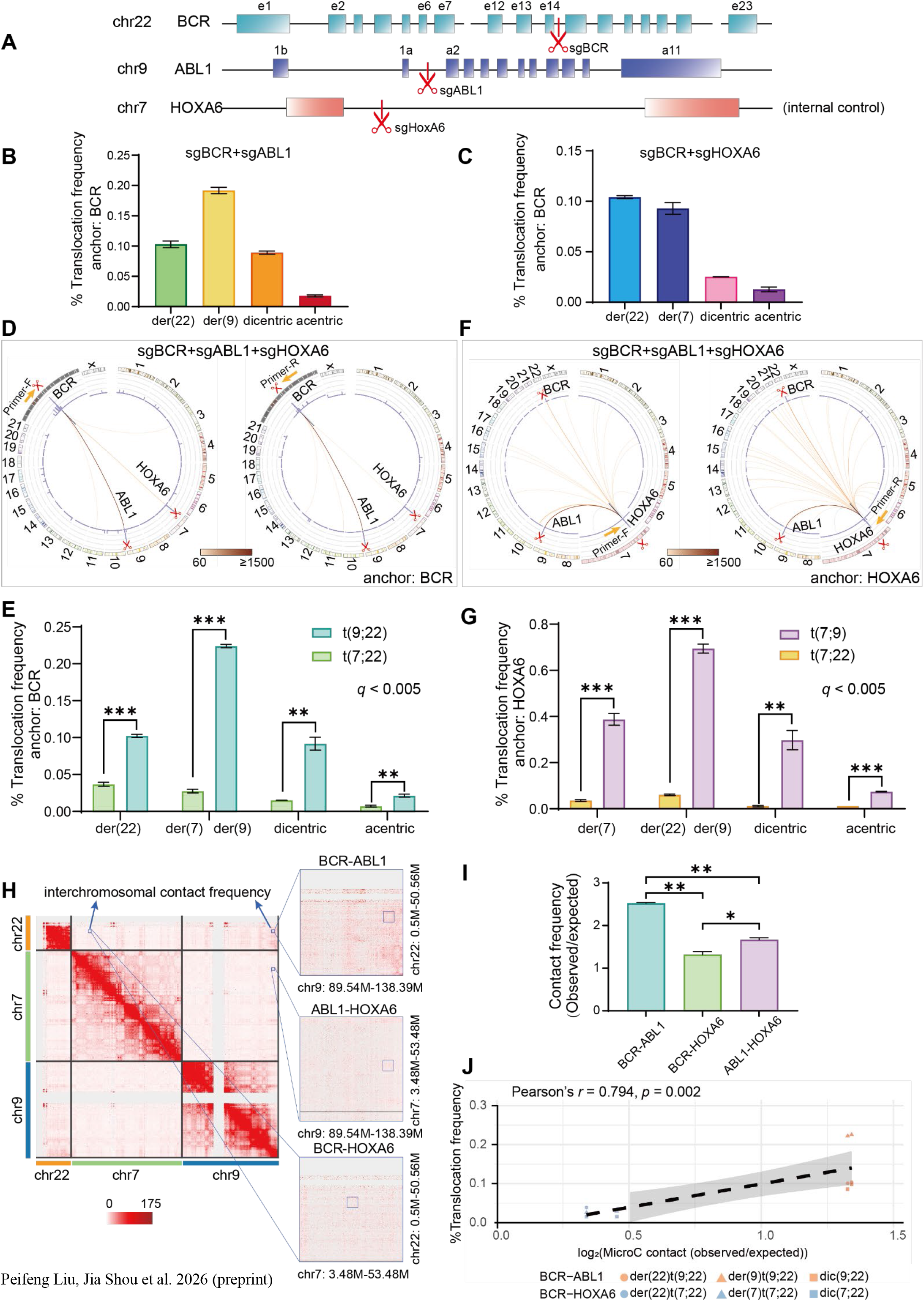
3D genome architecture affects chromosomal translocation. (A) Schematic of CRISPR/Cas9-mediated DSB induction at *BCR, ABL1*, and *HOXA6* loci for chromosomal translocation. (B, C) Translocation frequencies induced by dual sgRNAs at *BCR* and *ABL1* (B) or *BCR* and *HOXA6* (C) loci. (D-G) Chromosomal translocation analyses following simultaneous induction of DSBs with three sgRNAs. Translocation assays were anchored at *BCR* (D, E) or *HOXA6* (F, G). Circos plots show genome-wide translocation distributions; link color intensity reflects frequency; histograms denote translocation density across 500 bp bins (D, F). Quantifications of chromosomal translocations in D and F are shown in E and G, respectively. (H) Genome-wide Micro-C contact matrices (KR normalized, 500-kb resolution) with zoomed-in views (100-kb resolution) of spatial interactions among *BCR, ABL1*, and *HOXA6*. (I) Quantitative comparison of observed/expected Micro-C contact frequencies among the three loci. (J) Pearson correlation between translocation frequencies (anchored at *BCR*, excluding acentric fragments) and log2-transformed observed/expected Micro-C contact frequencies for three-DSB induction. The dotted line indicates linear regression fit. Data in B, C, E, G, and I are mean ± SD. *P* values were determined by two-tailed Student’s *t*-test with Benjamini-Hochberg correction (E, G) or two-tailed Student’s *t*-test (I).

### DSB end configuration determines translocation junctional pattern

We next investigated the role of DSB end configuration of Cas9 cleavage in chromosomal translocation. To this end, we quantified the relative contributions of blunt and staggered ends and found that translocation frequency correlates strongly with the proportion of staggered ends (Pearson *r* = 0.908, *p* < 0.001), suggesting that staggered DSB ends are intrinsically more prone to chromosomal translocation (Fig. 3A). We then profiled the *BCR-ABL1* junctions and found an indel landscape with prominent MMEJ-associated deletions and Polθ-mediated insertions, suggesting an MMEJ-driven repair mechanism in chromosomal translocation (Fig. 3B,C; Supplemental Fig. S4A-F).

**Fig 3.**
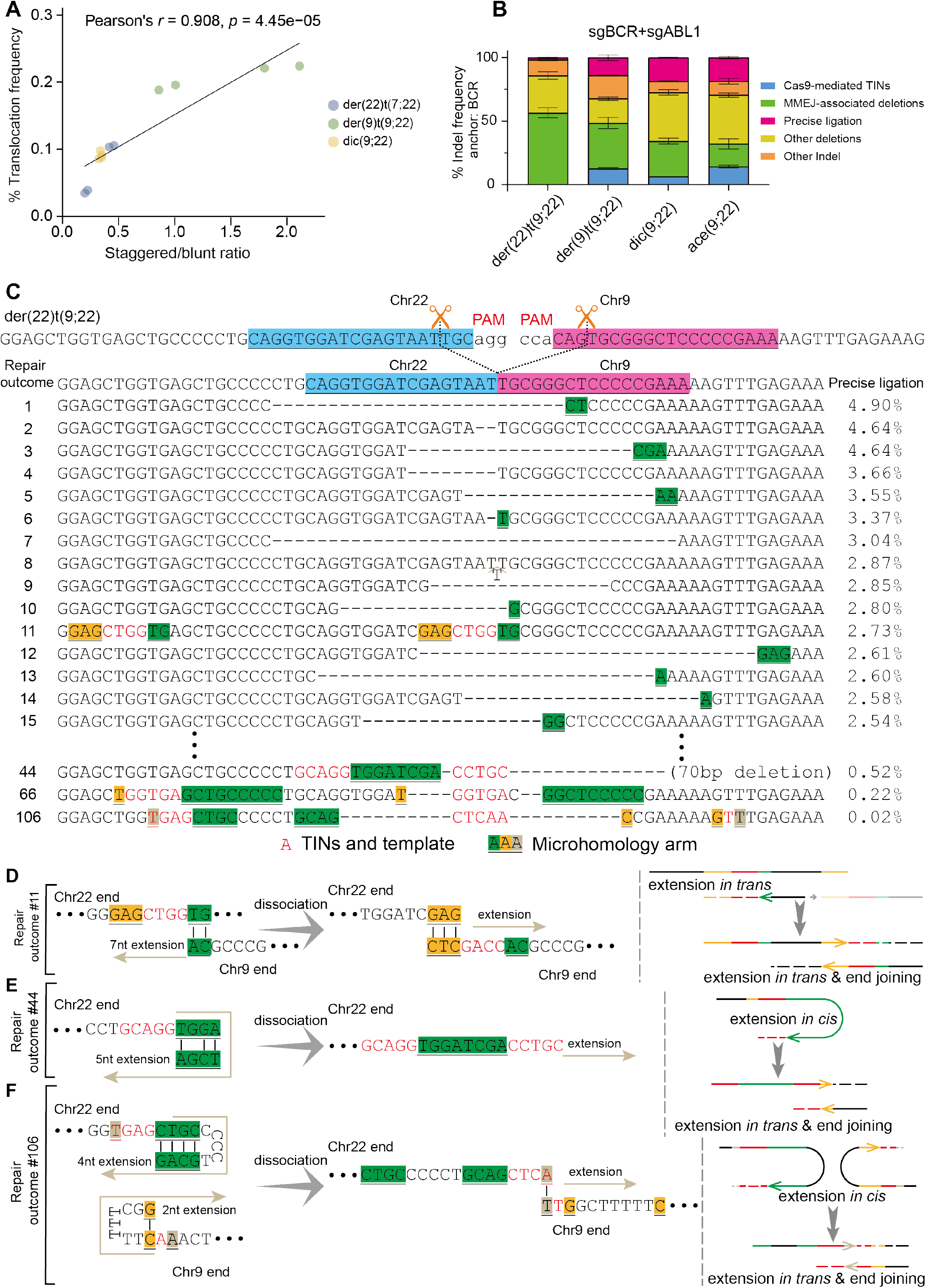
Cas9-induced DSB end configuration determines translocation junctional pattern. (A) Correlation between translocation frequency and the ratio of staggered versus blunt DSB ends induced by dual or triple sgRNAs. Acentric fragments, der(22)t(9;22), dic(7;22), and der(7)t(7;22) were excluded due to low detectability or inability to determine the staggered-to-blunt ratio. The solid line indicates the linear regression model. (B) Classification of repair outcomes at t(9;22) junctions into five categories. Data are mean ± SD. (C) Sequence alignment and classification of the top 15 repair outcomes at der(22)t(9;22) junctions. Deletions (dashes), templated insertions (red), and microhomology regions (highlighted and underlined) are indicated. (D-F) Mechanistic models of MMEJ-associated templated insertions (MMEJ-TINs) corresponding to repair outcomes in panel C: templated extension *in trans* (D, repair outcome #11), templated hairpin extension *in cis* (E, repair outcome #44), and combined templated extensions *in cis* on both ends (F, repair outcome #106).

In addition to canonical MMEJ-associated deletions, we identified a class of nucleotide insertions exhibiting high homology to sequences flanking the breakpoints. These insertions predominantly harbored multiple microhomology segments at the translocation junctions (Fig. 3C-F), and likely arise from Polθ-mediated extension *in cis* or *in trans*, followed by template dissociation and rejoining, generating MMEJ-associated templated insertions (MMEJ-TINs) (Kent et al. 2015). Representative examples within the der(22)t(9;22) junctions illustrate multi-step formation of such insertions (Fig. 3D-F). Mechanistically, DSB end resection exposes 3’ single-stranded DNA, enabling transient microhomology pairing, followed by Polθ-mediated templated extension and secondary pairing to bridge the breakage (Fig. 3D-F). Together, these observations suggest that Cas9-induced DSB end structures are crucial for chromosomal translocation.

### DISCOVER-seq reveals the *in-vivo* Cas9 staggered cleavage geometry during translocation

To directly characterize Cas9-generated DSB ends *in vivo*, we employed strand-resolved MRE11 ChIP-seq as established in the DISCOVER-seq (Wienert et al. 2019) to map Cas9 scissile sites at single-nucleotide resolution. Because MRE11 is rapidly recruited to DSB ends, the distributions of strand-specific reads accurately reflect Cas9 cleavage positions. We profiled three representative loci and resolved their cleavage sites at single nucleotide resolution (Fig. 4A-C). Strand-resolved profiling revealed pronounced Cas9 cleavage asymmetry. Specifically, cleavage on the complementary strand by the HNH domain was tightly enriched at the exactly -3 bp upstream of PAM, whereas cleavage on the noncomplementary strand by RuvC displayed positional variability, with peaks at -3, -4, or -5 bp upstream of PAM (Fig. 4A-C). Quantitative analyses confirmed that staggered DSB ends constitute a substantial fraction of Cas9-induced breaks *in vivo* (Fig. 4D-F). This heterogeneity indicates that Cas9 generates a mixture of blunt and staggered ends *in vivo* with 1-3 nt 5’ overhangs, which is consistent with our previous biochemical and genetic data (Shou et al. 2018).

**Fig 4.**
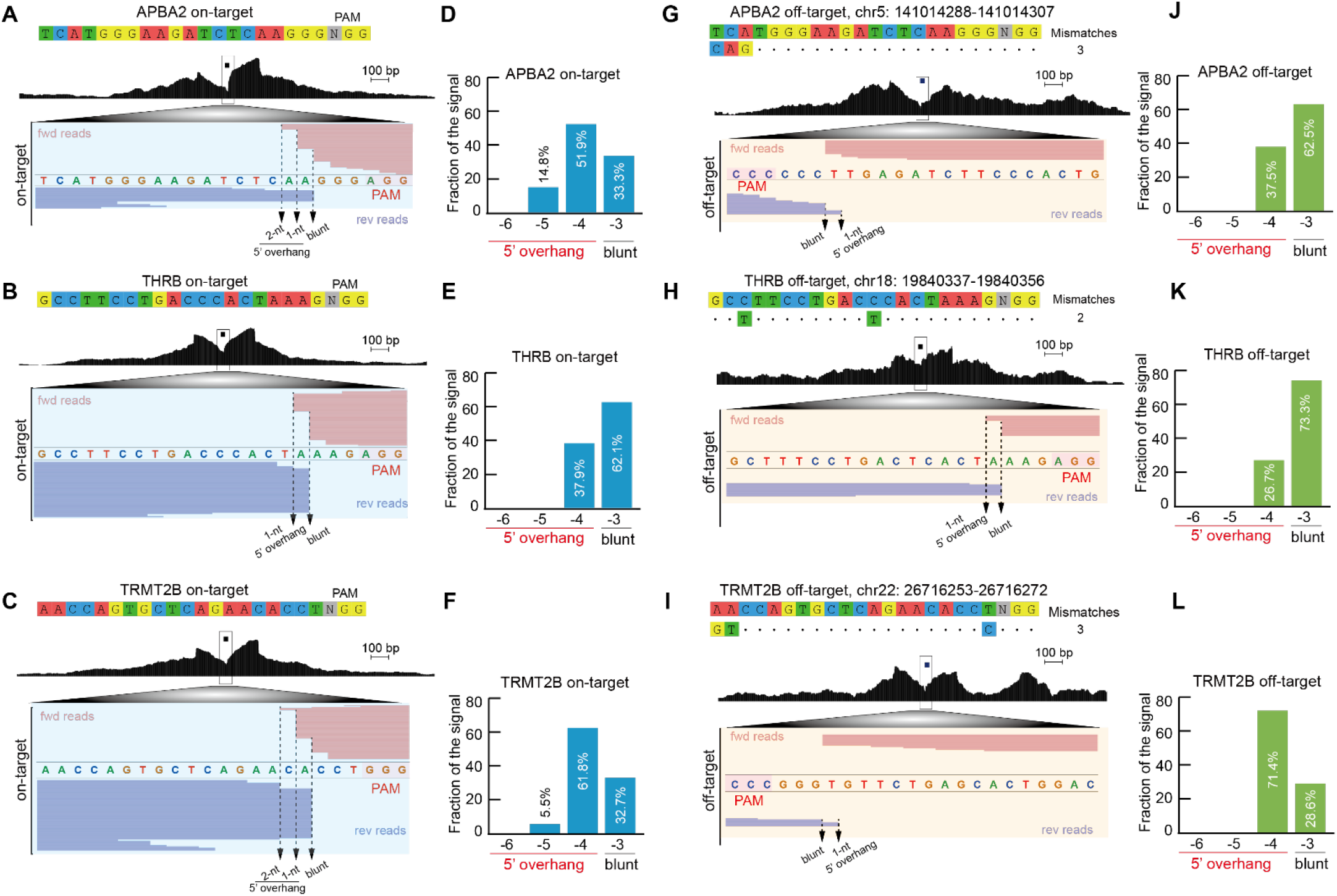
DISCOVER-seq resolves *in vivo* Cas9 staggered cleavage geometry at on-target and off-target sites. (A-C) Strand-specific DISCOVER-seq profiles at representative on-target loci (*APBA2, THRB*, and *TRMT2B*). Signals are shown across ±1 kb windows centered on the cleavage site. HNH domain cleavage (complementary strand) occurs 3 bp upstream of PAM; RuvC domain cleavage (non-complementary strand) exhibits positional heterogeneity. (D-F) Quantification of DNA end configurations at *APBA2, THRB*, and *TRMT2B* on-target sites. Noncomplementary strand cleavage by RuvC at position -3 produces blunt ends; cleavage at -4 to -6 generates staggered DSB ends with 5’ overhangs. (G-I) DISCOVER-seq profiles reveal asymmetric Cas9 cleavage signatures at off-target sites. (J-L) Quantification of DSB end configurations at off-target sites.

We then analyzed off-target sites corresponding to sgRNA targets of the *APBA2, THRB*, and *TRMT2B* genes, respectively. Remarkably, we also observed staggered cleavage at these off-target sites (Fig. 4G-I). Quantitative analyses revealed that the off-target staggered cleavage profiles are different from those of the on-target cleavages (Fig. 4J-L). Finally, we profiled 28 additional loci using DISCOVER-seq and found that staggered cleavage geometry is a general feature of Cas9 in both on-target and off-target activities (Supplemental Fig. S5).

### High-throughput GFP reporters for profiling chromosomal rearrangement junctions and Cas9 cleavage geometry

Based on the observed asymmetry in Cas9 cleavage, we hypothesized that cleavage geometry acts upstream of DNA repair pathway choice. Specifically, altering the position of the noncomplementary strand cleavage by the RuvC domain should reshape DSB end structures and influence repair outcomes. To this end, we established a gain-of-function dual-sgRNA reporter system that selectively captures repair outcomes of templated nucleotide insertions (Fig. 5A). Unlike previous reporters of EJ7-GFP (Bhargava et al. 2018), which primarily detect error-free blunt-end ligation, our system requires a precise +1 bp templated insertion to restore the GFP open reading frame (Supplemental Fig. S6A,B). Thus, GFP recovery directly reflects the fill-in synthesis of 1-nt 5’ overhangs.

**Fig 5.**
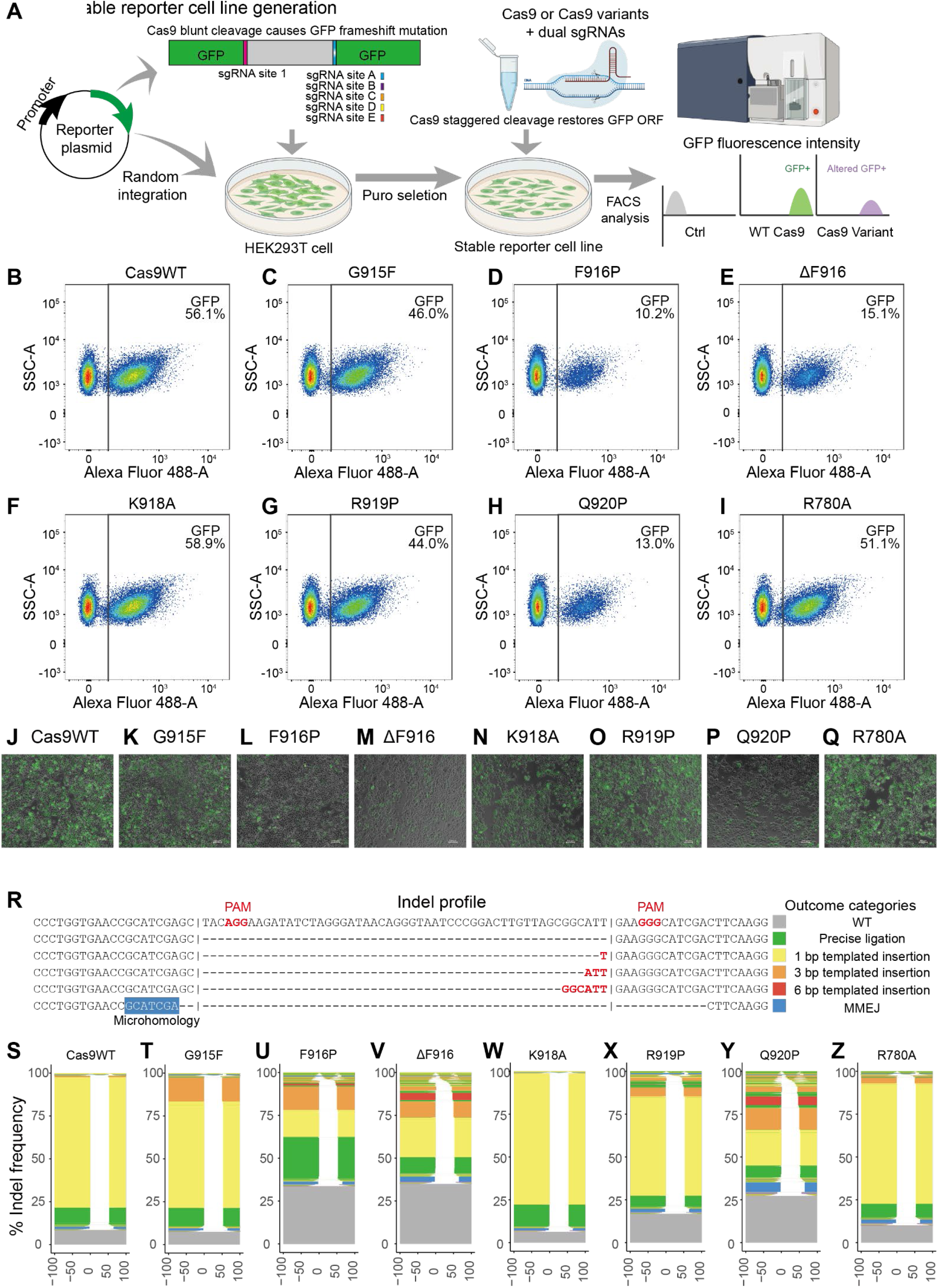
High-throughput reporter profiling of engineered Cas9 cleavage geometry. (A) Schematic of the GFP-based dual-sgRNA reporter system. Functional GFP restoration specifically requires a precise +1 bp templated insertion indicative of staggered cleavage. (B-I) Flow cytometric quantification of GFP-expressing HEK293T cells edited with of Cas9WT (B) or engineered Cas9 variants of G915F (C), F916P (D), ΔF916 (E), K918A (F), R919P (G), Q920P (H), and R780A (I). (J-Q) Fluorescence microscopy of GFP-expressing HEK293T cells with editors of Cas9WT or engineered Cas9 variants. (R-Z) Sequence alignments of five types of repair outcomes (R) and their normalized indel frequencies (S-Z) at reporter junctions with editors of Cas9WT (S) or engineered Cas9 variants of G915F (T), F916P (U), ΔF916 (V), K918A (W), R919P (X), Q920P (Y), and R780A (Z). Note the shifted 5’ overhang lengths and increased frequencies of larger size of templated insertion for F916P (U), ΔF916 (V), and Q920P (Y).

Using this system, we comprehensively tested engineered Cas9 variants including G915F, F916P, ΔF916, K918A, R919P, Q920P, and R780A (Fig. 5B-Q). Flow cytometric analyses revealed that wild-type Cas9 exhibited robust GFP recovery (∼56%) (Fig. 5B), consistent with frequent generation of 1-nt 5’ overhangs. Notably, F916P, ΔF916, and Q920P produced markedly altered fluorescence profiles (Fig. 5D,E,H). Live-cell imaging confirmed the significant decrease of GFP fluorescence for these three variants (Fig. 5L,M,P). Deep sequencing revealed that they generate distinct indel profiles (Fig. 5R-Z; Supplemental Fig. S6C). Specifically, whereas wild-type Cas9 predominantly produced +1 bp insertions corresponding to RuvC cleavage at -4 bp upstream of PAM, these variants shifted cleavage further upstream at -6 or -9 bp of PAM, yielding 3 or 6 bp templated insertions (Fig. 5U,V,Y). In particular, F916P, ΔF916, and Q920P generate remarkable 1.06%, 3.93%, and 4.74% of 6-bp templated insertions, respectively, as highlighted by the red color (Fig. 5U,V,Y). Finally, clustering analyses of junctional indel patterns revealed that these variants form a distinct group from the other five ones (Supplemental Fig. S6D). These data suggest that chromosomal rearrangement junctions and Cas9 cleavage geometry can be altered by Cas9 protein engineering.

## Discussion

Accumulating evidence suggests that chromosomal translocation formation is shaped by both nuclear spatial organization and DNA-repair pathway choice (Mani and Chinnaiyan 2010; Roukos and Misteli 2014). Initial studies using I-SceI induced cleavage showed that three-dimensional (3D) proximity between genomic loci is a key determinant of translocation frequency(Zhang et al. 2012). In addition, recurrent translocation partners in clinical samples often display elevated contact frequencies prior to breakage induction and that translocation frequency correlates strongly with pre-existing chromatin interactions (Engreitz et al. 2012). We sought to investigate the relationship between frequencies of chromosomal rearrangements and the size of the intervening DNA fragments using the CRISPR/Cas9 system programmed with dual sgRNAs (Guo et al. 2015; Li et al. 2015; Shou et al. 2018; Shi et al. 2019; Mehryar et al. 2023). Intriguingly, we observed frequent chromosomal rearrangements between very distant genomic sites within single chromosomes, suggesting that spatial but not linear distance plays a role in the end-joining of DSBs (Supplemental Table S2). We reasoned that erroneous end-joining between non-homologous chromosomes would be similar to that between genomic sites very far apart within the same chromosome. Indeed, we observed chromosomal translocations in 77 out of 90 cases involving Cas9-induced DSBs at random genomic sites of different chromosomes (Supplemental Table S1). We report for the first time that junctional indel patterns at chromosomal translocations are determined by the relative orientation and geometry of Cas9-induced staggered cleavages. In particular, Cas9 does not exclusively produce blunt DSBs *in vivo* but instead frequently generates short 5’ overhangs during chromosomal translocation. This intrinsic bias provides a mechanistic explanation for the templated insertions observed at translocation junctions (Fig. 1), as the 5’ overhang nucleotides can be directly incorporated during end-joining of DSB repair. Furthermore, strand-resolved MRE11 ChIP-seq shows the flexibility of Cas9 cleavage on the non-complementary strand, independently confirming the staggered end geometry inferred from junctional sequence analyses.

The configuration of DSB ends and the choice of repair pathway influence the sequence architecture of translocation junctions. Both classical non-homologous end joining (c-NHEJ) and alternative end joining (alt-EJ/MMEJ) contribute to translocation formation, with their relative engagement determined by DSB end structure and cellular context (Ghezraoui et al. 2014; Chang et al. 2016; Bothmer et al. 2017). However, the impact of DSB end configuration on translocation frequency and junctional structure remains poorly characterized in CRISPR-induced models. Although Cas9 has been widely assumed to generate blunt DSB ends, recent studies indicates that the RuvC domain exhibits cleavage flexibility on the non-complementary strand, often producing staggered ends with 5’ overhangs(Shou et al. 2018; Shi et al. 2019; Chauhan et al. 2023; Chauhan et al. 2025). Rather than functioning solely as a blunt-end endonuclease with 3’-5’ exonuclease activity, Cas9 frequently generates 5’ overhangs due to variability in RuvC cleavages of the non-complementary strand (Jinek et al. 2012; Shou et al. 2018; Chauhan et al. 2025). Importantly, because the formation of a translocation junction disrupts the original sgRNA target sequences, the 3-bp and 6-bp templated insertions observed at chromosomal translocation junctions cannot be explained by iterative ligation and recleavage (Figs. 1L and 5R). This reinforced the conclusion that Cas9 intrinsically produces staggered DSB ends and directly challenged the recent model attributing multi-nucleotide templated insertions to repeated blunt-end cutting and 1-bp insertion (de Alba et al. 2025).

Nuclear organization establishes the likelihood of interactions between DSB ends, as evidenced by the strong correlation between Micro-C contact frequencies and translocation rates, indicating that recurrent oncogenic rearrangements such as *BCR-ABL1* are structurally preconfigured by genome architecture prior to any selective pressure (Fig. 2). However, spatial proximity alone does not fully explain the efficiency or sequence features of translocation junctions. DSB end configuration provides an additional layer of regulation by determining how encounters are resolved into stable rearrangements. In particular, staggered ends exhibit increased translocation propensity compared with blunt ends (Fig. 3), consistent with recent studies that DSB ends with 5’ overhangs impact DNA recombination, leading to chromosomal translocations (So and Martin 2019). In addition, junctional signatures support a hierarchical repair process in which c-NHEJ-mediated fill-in precedes MMEJ, linking DSB end geometry to repair pathway choice. These observations suggest that end configuration itself is a key determinant of repair outcomes. Taken together, we propose a unified model in which genome architecture constrains the probability of DSB-end encounters, while DSB end configuration affects ligation compatibility (Supplemental Fig. S6E). Further investigation will be required to elucidate how CRISPR-induced end heterogeneity cooperates with nuclear architecture to shape the mechanisms underlying chromosomal translocation formation.

## Materials and methods

### Cell lines and culture conditions

HEK293T cells (human embryonic kidney 293T; ATCC CRL-3216) were maintained in Dulbecco’s modified Eagle’s medium (DMEM; Gibco, C11995500BT) supplemented with 10% (v/v) fetal bovine serum (FBS; Gibco, A5256701) and 1% (v/v) penicillin-streptomycin (PS; Gibco, 10378-016). Cells were cultured at 37°C in a humidified incubator containing 5% CO_2_. All cell lines were regularly tested for mycoplasma contamination by PCR and confirmed to be mycoplasma-free.

### SgRNA design and plasmid construction

The sgRNA target sequences were designed using CRISPOR (https://crispor.gi.ucsc.edu) to maximize predicted on-target specificity while minimizing off-target activity. Candidate sgRNAs were ranked according to their specificity scores, and those with the highest specificity and lowest predicted off-target burden were selected. For the dual-sgRNA reporter system, the inDelphi algorithm (Shen et al. 2018) was employed to predict indel profiles at the intended cleavage sites. All sgRNA sequences are listed in Supplemental Table S2.

To generate sgRNA expression plasmids, a pair of complementary oligonucleotides containing the targeting sequence (Supplemental Table S2) were annealed and ligated into the BsaI-linearized pGL3-U6-sgRNA vector (provided by Prof. X. Huang, ShanghaiTech University) downstream of the U6 promoter.

For the construction of the dual-sgRNA reporter, three fragments corresponding to the N-terminal GFP half, the C-terminal GFP half, and the dual-sgRNA cassette containing an internal linker sequence were amplified by PCR using primers listed in Supplemental Table S2. The fragments were assembled by overlap extension PCR, and the resulting product was digested and ligated into the linearized pLVX vector.

To generate tandem sgRNA constructs for the dual sgRNA reporter, two copies of the sgRNA expression cassette, each consisting of a human U6 promoter, one split-GFP-targeting sgRNA sequence, and the sgRNA scaffold, were inserted into the lentiCRISPR plasmid (Addgene, #52961). The Cas9 coding sequence was subsequently removed from the modified backbone, and the puromycin resistance cassette was replaced with a hygromycin resistance cassette to obtain the final sgRNA expression construct.

All plasmids were transformed into competent *Escherichia coli* stbl3 cells, and plasmid DNA was isolated from individual colonies using a miniprep kit (Axygen, AP-MN-P-250). The sequences and orientations of all sgRNA constructs were confirmed by Sanger sequencing.

### Transfection and cell harvesting

One day prior to transfection, HEK293T cells were seeded at a density of 8 × 10^5^ cells per well in six-well plates to achieve 60-80% confluency at the time of transfection. For each well, a plasmid mixture containing the SpCas9-GFP expression vector and the sgRNA expression plasmid at a mass ratio of 2:3 was prepared. Transfection was performed using Lipofectamine 3000 (Thermo Fisher, L3000015) according to the manufacturer’s instructions. Cells were incubated for 48 h, and the culture medium was replaced 24 h post-transfection.

To isolate successfully transfected cells, GFP-positive populations were sorted by fluorescence-activated cell sorting (FACS) using a BD FACS instrument. Prior to sorting, cells were detached by trypsinization, washed twice with PBS, and resuspended in PBS containing 2% FBS. Sorted GFP-positive cells were collected into complete medium and cultured for an additional 7 days to allow recovery and clonal expansion. Genomic DNA was then extracted using a standard phenol-chloroform method, quantified using a NanoDrop 2000 instrument (Thermo Fisher), and stored at - 20°C for subsequent analyses.

### PCR detection and junction characterization of chromosomal translocations

Chromosomal translocations were detected by PCR using optimized primer pairs flanking the predicted breakpoint regions. Primer combinations were designed based on the relative orientations and chromosomal locations of the target loci. When the two loci resided on different chromosomal arms (p and q), amplification was performed using either forward-forward or reverse-reverse primer pairs. In contrast, when both loci were located on the same arm, a forward primer from one locus was paired with a reverse primer from the other to enable junction-specific amplification. PCR products were resolved by agarose gel electrophoresis, excised, and purified, followed by cloning and sequencing (TSV-007VS, Tsingke). Individual colonies were selected and subjected to Sanger sequencing to confirm the presence of translocation events and to characterize breakpoint junction sequences at single-nucleotide resolution.

### Genome-wide detection of chromosomal translocation and next-generation sequencing

Translocation junctional libraries were constructed as previously described (Yin et al. 2019; Liu et al. 2021) with minor modifications. Briefly, approximately 20 μg of genomic DNA isolated from edited cells was fragmented by sonication to generate fragments predominantly ranging from 300 to 1000 bp. The fragmented DNA was subjected to primer extension using a biotinylated bait primer (Supplemental Table S3) targeting a region located approximately 100 bp upstream of the cleavage site. A linear amplification step is applied in 100 μl of PCR system for a total of 80 cycles. Extension products were denatured at 95°C for 5 min and captured using streptavidin-coated magnetic beads (Thermo Fisher, 65001). Captured DNA was ligated on-bead overnight at 16°C to a pre-annealed double-stranded adapter. Bead-bound libraries were then amplified by nested PCR using indexed primers containing Illumina P5/P7 adaptor sequences (Supplemental Table S2). Final libraries were sequenced on an Illumina NovaSeq X Plus platform in 150 bp paired-end mode.

Raw sequencing reads were preprocessed by adapter trimming with Cutadapt (v4.9) (Martin 2011) followed by quality filtering (Phred score ≥ 20). To minimize non-specific alignments, reads were retained only when concordantly mapped to both the primer sequence and at least 5 bp of the immediately downstream genomic sequence, ensuring that each read originated precisely from the primer binding site. PCR duplicates were removed using FastUniq (Xu et al. 2012). Following the PEM-Q pipeline (Liu et al. 2021), R1 reads were aligned to the human reference genome hg19 using BWA-MEM (v0.7.17) (Li and Durbin 2009) with default parameters. Chimeric reads were identified through SA tags and classified into different chromosomal translocations. Translocation frequencies were normalized to reads per million reads (RPM).

### Characterization of breakpoint junction features

To characterize sequence features at DSB repair junctions, locus-specific primers flanking the *BCR, ABL1, HOXA6* and dual sgRNA GFP reporter breakpoint regions were designed using Primer-BLAST (https://www.ncbi.nlm.nih.gov/tools/primer-blast) (Supplemental Table S2). Primer pairs were validated *in silico* against the hg19 reference genome and further tested by conventional PCR. The junctions at the *BCR, ABL1* and *HOXA6* cleavage sites were characterized by conventional PCR amplification using locus-specific primer pairs, followed by third-generation sequencing. For the dual sgRNA GFP reporter, junctional indel patterns were characterized by PCR-based library preparation using indexed primers containing 5’ Illumina P5/P7 adaptor sequences and 3’ locus-specific primers, followed by next-generation sequencing on an Illumina NovaSeq X Plus platform in 150 bp paired-end mode.

All high-throughput sequencing data regarding breakpoint junctional features were analyzed using a standardized analytical pipeline. Repair outcomes were classified according to junctional indel patterns (templated insertion, deletion, precise ligation, indels) as previously described (Mehryar et al. 2023). Briefly, raw reads were first subjected to quality control to remove nonspecific amplification products, low-quality sequences, adapter contamination, and excessively short fragments. High-quality chimeric reads were aligned to references reconstructed from the predicted junction-associated segments. To identify templated insertions, each reference segment was extended with sequences from the opposite side of the cleavage site. Because these sequences are absent from the final translocation reads, templated insertions are often recognized as random insertions. Our reference design and alignment strategy effectively prevents such incorrect recognition. Segment-level junctional indel patterns were subsequently integrated to generate read-level indel patterns at single-nucleotide resolution. Junctional features, including insertion length, deletion size, templated insertion origin, and microhomology usage, were further quantified for subsequent statistical analyses. This analytical pipeline enabled quantitative determination of the type and frequency of indel events at individual junctions. Frequencies of each junctional indel pattern were normalized to total reads per junction.

### Third-generation sequencing

Junction-spanning amplicons covering *BCR, ABL1* and *HOXA6* cleavage sites were generated by locus-specific primers pairing using Phanta Max Super-Fidelity DNA Polymerase (Vazyme, P505-d1) under the following conditions: 95°C for 3 min; 35 cycles of 95°C for 15 s, 60°C for 15 s, and 72°C for 1 min; followed by a final extension at 72°C for 5 min. PCR products were subjected to third-generation nanopore sequencing (Oxford Nanopore Technologies). Library preparation and sequencing were performed following standard ONT protocols. ONT sequencing enables direct, long-read characterization of PCR amplicons without the need for amplification during sequencing, allowing full-length reads of target fragments. Data processing was performed identically to sequencing analyses of breakpoint junction features described above. Reads were aligned to the references, and indel patterns relative to the reference were identified and statistically analyzed for estimation of sgRNA-mediated cleavage efficiency at each target locus.

### Micro-C data processing and visualization

Publicly available Micro-C data generated from HEK293T cells (Shanel et al. (Shanel et al. 2025); GEO accession: GSE278978) were downloaded from the GEO database. AllValidPairs files were converted to .hic format using the hicpro2juicebox script included in HiC-Pro (v2.11.1) (Servant et al. 2015). The resulting files were imported into Juicer Tools (v2.20.00) (Durand et al. 2016b) for matrix extraction and Knight-Ruiz (KR) normalization. Genome-wide contact matrices were visualized at multiple resolutions using Juicebox (v2.15) (Durand et al. 2016a) and displayed as observed/expected ratios.

### DISCOVER-seq library construction and data analyses

DISCOVER-seq libraries were prepared as previously described (Wienert et al. 2019) with minor modifications. Briefly, one day before transfection, HEK293T cells were seeded at 6 × 10^6^ cells per 10-cm dish. SpCas9 and sgRNA expression plasmids were co-transfected using Lipofectamine 3000 (Thermo Fisher, L3000015) according to the manufacturer’s instructions. At 10 h post-transfection, the DNA-PK inhibitor KU-0060648 (SML1257) was added to the medium at a final concentration of 0.5 μM to enrich MRE11 occupancy at Cas9-induced DSB sites by suppressing NHEJ-mediated end joining. After 48 h of drug treatment, cells were harvested and cross-linked with 1% formaldehyde (Thermo Fisher, 28906) for 10 min at room temperature with gentle rotation. Cross-linking was quenched by addition of glycine to a final concentration of 125 mM, followed by incubation for 10 min at room temperature. Cells were pelleted at 1000 × g for 5 min at 4°C and washed twice with ice-cold PBS containing 1 × protease inhibitor cocktail (Roche, 04693116001).

Cross-linked cells were lysed, and nuclei were pelleted by centrifugation. Chromatin was fragmented by sonication to yield DNA fragments of approximately 150-500 bp. Lysates were pre-cleared with protein A/G magnetic beads (Thermo Fisher, 88802), and 5% of the material was reserved as input. The remaining lysate was incubated overnight at 4°C with anti-MRE11 antibody (NB100-142, NOVUS) under gentle rotation. The following day, protein A/G magnetic beads were added and incubated for 3 h at 4°C with gentle rotation to capture antibody-chromatin complexes. The beads were sequentially washed with ChIP buffers and eluted from the beads with elution buffer. Cross-links were reversed by overnight incubation at 65°C with shaking at 1000 rpm. Following RNase A (Thermo Fisher, EN0531) and Proteinase K (Thermo Fisher, EO0491) treatment, ChIP DNA was purified using the MinElute Gel Extraction Kit (Qiagen, 28606) and quantified by Qubit (Thermo Fisher).

Sequencing libraries from ChIP DNA and matched input controls were prepared using the Universal DNA Library Prep Kit for Illumina (Vazyme, ND607) according to the manufacturer’s protocol, including end repair, 3′-dA tailing, dual-index adapter ligation, and PCR amplification. Final libraries were size-selected to 200-500 bp using AMPure XP beads (BECKMAN COULTER, A63881) and sequenced on an Illumina NovaSeq X Plus platform in 150 bp paired-end mode.

Raw sequencing reads were processed using Trim Galore (v0.6.7) for adapter removal and low-quality trimming (Phred score < 20). Filtered reads were aligned to hg19 using Bowtie2 (v2.5.4) (Langmead and Salzberg 2012). SAM files were converted to sorted and indexed BAM files using SAMtools (v1.19.2) (Li et al. 2009). PCR duplicates were marked and removed using Picard MarkDuplicates (v3.1.1) with default settings. Mapping rate, duplication rate, and library complexity were evaluated simultaneously.

Genome-wide normalized coverage tracks were generated using bamCoverage in deepTools (v3.5.5) (Ramírez et al. 2014) with reads per kilobase per million mapped reads (RPKM) normalization. Genome-wide MRE11 enrichment was visualized using the R package karyoploteR (Gel and Serra 2017), with on-target and off-target loci annotated.

Genome-wide off-target sites for each sgRNA were predicted using Cas-OFFinder (v2.4) (Bae et al. 2014) against hg19 reference genome, allowing up to 5 bp mismatches relative to the on-target protospacer sequence. For the 31 sgRNAs used in this study, 103,397 potential off-target sites were identified. DISCOVER-seq signals were extracted from all on-target and predicted off-target loci. Sites showing read coverage on both plus-strand and minus-strand read initiation ends were retained, yielding approximately 4,138 high-confidence candidate sites. These loci were ranked by read depth and sequentially inspected in IGV (v2.19.5) (Thorvaldsdóttir et al. 2013).

### Establishment of stable cell lines for dual sgRNA reporter

Recombinant lentiviral particles were generated by transient co-transfection of HEK293T cells. One day before transfection, cells were seeded at 8 × 10^5^ cells per well in six-well plates containing antibiotic-free complete DMEM. For each well, a three-plasmid packaging system was used, consisting of the pLVX-dual sgRNA reporter transfer plasmid, psPAX2 (Addgene, 12260), and pMD2.G (Addgene, 12259) at a molar ratio of 4:3:1. Transfection was performed using Lipofectamine 3000 (Thermo Fisher, L3000015) according to the manufacturer’s instructions. At 8-12 h post-transfection, the medium was replaced with fresh DMEM supplemented with 20% FBS to enhance cell viability and maximize viral production.

Virus-containing supernatants were collected at 36 h and 60 h post-transfection. Each harvest was centrifuged at 1000 × *g* for 5 min at 4°C to remove cellular debris and filtered through a 0.45 μm PVDF membrane (Millex, SLHVR33RB). Supernatants were concentrated using PEG-8000. After thorough mixing and overnight incubation at 4°C, samples were centrifuged at 4,000 × *g* for 20 min at 4°C, and the supernatant was discarded. Viral pellets were resuspended in an appropriate volume of fresh DMEM and stored at -80°C.

Target HEK293T cells were seeded at 1-2 × 10^6^ cells per well in six-well plates containing 2 ml complete DMEM. Before transduction, the medium was replaced with 2.5 ml antibiotic-free DMEM supplemented with 10% FBS and 8 μg/ml polybrene (Beyotime, C0351-1ml). Concentrated viral supernatant (250 μl) was added dropwise to each well. After 48 h, the medium was replaced with selection medium containing 2.5 μg/ml puromycin (Solarbio, P8230).

## Statistical analyses

All quantitative data are presented as mean ± standard deviation (SD) from at least two independent biological replicates unless otherwise indicated. Statistical significance for pairwise comparisons was determined using an unpaired two-tailed Student’s *t*-test. For experiments involving more than three groups, one-way analysis of variance (ANOVA) was performed. When more than four independent comparisons were conducted within a study, *p*-values were adjusted using the Benjamini-Hochberg false discovery rate (FDR) method to control type I error. All analyses were performed using GraphPad Prism (v9.5.1) and R (v4.5.2). Statistical significance was defined as * *p* ≤ 0.05, ** *p* ≤ 0.01, *** *p* ≤ 0.001, and NS (not significant). Pearson correlation coefficient was used to assess linear associations between variables, and linear regression lines were fitted to scatter plots to illustrate overall trends.

## Data visualization

Genomic tracks were visualized using the UCSC Genome Browser (Casper et al. 2026) or Integrative Genomics Viewer (IGV, v2.19.5) (Thorvaldsdóttir et al. 2013). Chromosomal rearrangement diagrams and interchromosomal translocation maps were generated using Circos (v0.69) (Krzywinski et al. 2009). Statistical graphics were produced in R (v4.5.2) using ggplot2 (v4.0.2) and GraphPad Prism (v9.5.1).

## Acknowledgments

We are very grateful to Dr. Hongbo Gao and Yanwen Xue from the School of Agriculture and Biology of SJTU for the help of FACS, Juan Yang from the State Key Laboratory of Oncogenes & Related Genes of SJTU for the help of many basic laboratory instruments. We are also grateful for advice on bioinformatics from Dr. Jingwei Li and all members of our laboratory for discussion. This study was supported by grants from the National Natural Science Foundation of China (32330016 to Q.W. and 32000425 to J.S.), the National Key R&D Program of China (2022YFC3400200 to Q.W.), the Shanghai Natural Science Foundation (25ZR1402243 to J.S.), and the Postdoctoral Fellowship Program and China Postdoctoral Science Foundation (BX20200217 and 2020M681283 to J.S.).

## Author contributions

Conceptualization was by Q.W. Methodology, formal analysis, and investigation were by P.L. and J.S. Original draft was by P.L. and J.S. Editing was by Q.W. All authors read and approved the final manuscript.

