## Supplemental Figures for "Double-strand break end configuration and 3D genome architecture are crucial for chromosomal translocation"

**The PDF file includes:**

**Supplementary Figures S1 to S6**

**Fig S1.** PCR detection and Sanger sequencing confirmation of chromosomal translocations.

**Fig S2.** Translocation landscapes and junctional features of unbalanced rearrangements.

**Fig S3.** Translocation mapping and spatial correlations programed by dual or triple sgRNAs.

**Fig S4.** Representative junction features of Cas9-induced and Polθ-mediated templated insertions at translocation junctions.

**Fig S5.** DISCOVER-seq mapping of Cas9 cleavages at on-target and off-target sites.

**Fig S6.** Engineered Cas9 variants modulate end geometry and repair outcomes.

**Captions for Tables S1 to S3**

**Table S1.** Chromosomal translocations assayed by PCR.

**Table S2.** Chromosomal rearrangements within single chromosomes assayed by PCR.

**Table S3.** Oligonucleotides used in the study.

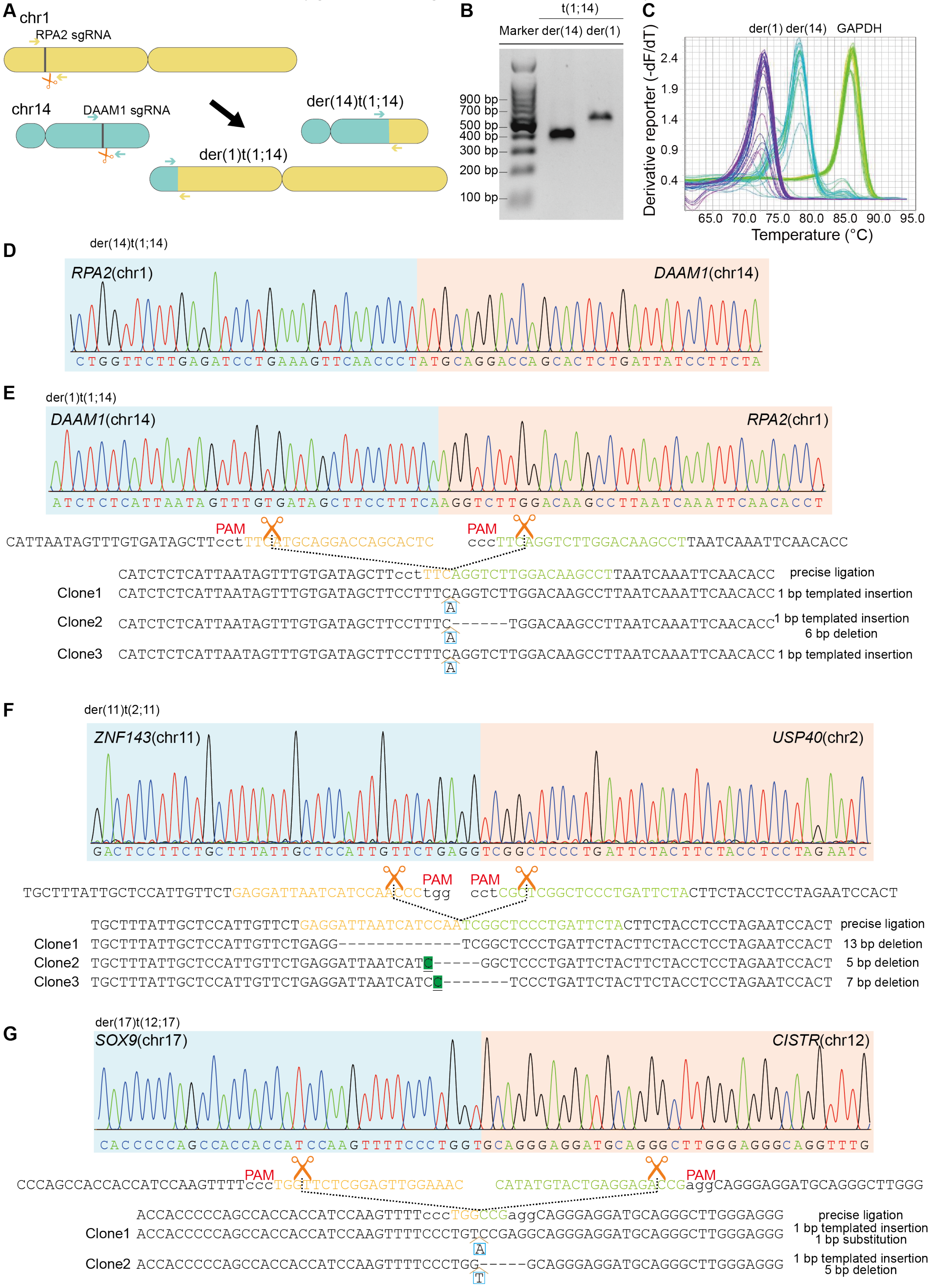
A
chr1
RPA2 sgRNA
chr14
DAAM1 sgRNA
der(14)t(1;14)
der(1)t(1;14)
B
t(1;14)
Marker der(14) der(1)
900 bp
700 bp
500 bp
400 bp
300 bp
200 bp
100 bp
C
der(1) der(14) GAPDH
Derivative reporter (-dF/dT)
2.4
1.9
1.4
0.9
0.4
65.0 70.0 75.0 80.0 85.0 90.0 95.0
Temperature (°C)
D
der(14)t(1;14)
RPA2(chr1)
DAAM1(chr14)
CTGGTTCTTGAGATCCTGAAAGTTCAACCCTATGCAGGACCAGCACTCTGATTATCCTTCTA
E
der(1)t(1;14)
DAAM1(chr14)
RPA2(chr1)
ATCTCTCATTAATAGTTTGTGATAGCTTCCTTTCAAGGTCTTGGACAAGCCTTAATCAAATTCAACACCT
PAM
PAM
CATTAATAGTTTGTGATAGCTTcctTTCATGCAGGACCAGCACTC
cccTTCAGGTCTTGGACAAGCCTTAATCAAATTCAACACC
CATCTCTCATTAATAGTTTGTGATAGCTTcctTTCAGGTCTTGGACAAGCCTTAATCAAATTCAACACC precise ligation
Clone1 CATCTCTCATTAATAGTTTGTGATAGCTTCCTTTCAGGTCTTGGACAAGCCTTAATCAAATTCAACACC 1 bp templated insertion
A
Clone2 CATCTCTCATTAATAGTTTGTGATAGCTTCCTTTC------TGGACAAGCCTTAATCAAATTCAACACC 1 bp templated insertion 6 bp deletion
A
Clone3 CATCTCTCATTAATAGTTTGTGATAGCTTCCTTTCAGGTCTTGGACAAGCCTTAATCAAATTCAACACC 1 bp templated insertion
A
F
der(11)t(2;11)
ZNF143(chr11)
USP40(chr2)
GACTCCTTCTGCTTTATTGCTCCATTGTTCTGAGGTCGGCTCCCTGATTCTACTTCTACCTCCTAGAATC
PAM
PAM
TGCTTTATTGCTCCATTGTTCTGAGGATTAATCATCCAATCCtgg
cctCGCTCGGCTCCCTGATTCTACTTCTACCTCCTAGAATCCACT
TGCTTTATTGCTCCATTGTTCTGAGGATTAATCATCCAATCGGCTCCCTGATTCTACTTCTACCTCCTAGAATCCACT precise ligation
Clone1 TGCTTTATTGCTCCATTGTTCTGAGG-------------TCGGCTCCCTGATTCTACTTCTACCTCCTAGAATCCACT 13 bp deletion
Clone2 TGCTTTATTGCTCCATTGTTCTGAGGATTAATCATC-----GGCTCCCTGATTCTACTTCTACCTCCTAGAATCCACT 5 bp deletion
Clone3 TGCTTTATTGCTCCATTGTTCTGAGGATTAATCATCC-------TCCCTGATTCTACTTCTACCTCCTAGAATCCACT 7 bp deletion
G
der(17)t(12;17)
SOX9(chr17)
CISTR(chr12)
CACCCCCAGCCACCACCATCCAAGTTTTCCCTGGTGCAGGGAGGATGCAGGGCTTGGGAGGGCAGGTTTG
PAM
PAM
CCCAGCCACCACCATCCAAGTTTTcccTGGTTCTCGGAGTTGGAAAC
CATATGTACTGAGGAGACCGaggCAGGGAGGATGCAGGGCTTGGG
ACCACCCCCAGCCACCACCATCCAAGTTTTcccTGGCCGaggCAGGGAGGATGCAGGGCTTGGGAGGG precise ligation
Clone1 ACCACCCCCAGCCACCACCATCCAAGTTTTCCCTGTCCGAGGCAGGGAGGATGCAGGGCTTGGGAGGG 1 bp templated insertion 1 bp substitution
A
Clone2 ACCACCCCCAGCCACCACCATCCAAGTTTTCCCTGG-----GCAGGGAGGATGCAGGGCTTGGGAGGG 1 bp templated insertion 5 bp deletion
T

**Fig S1. PCR detection and Sanger sequencing confirmation of chromosomal translocations.** (A) Dual-sgRNA CRISPR/Cas9 system for chromosomal translocation. Simultaneous DSB inductions at *RPA2* (chr1) and *DAAM1* (chr14) with Cas9 programmed by dual sgRNAs generate reciprocal translocations: der(1)t(1;14) and der(14)t(1;14). (B) Gel electrophoresis shows the PCR products of der(1)t(1;14) and der(14)t(1;14). (C) Quantitative PCR shows the melting curves of der(1)t(1;14), der(14)t(1;14), and the internal control (*GAPDH*). (D,E) Sanger sequencing chromatograms show the translocation junctions of der(14)t(1;14) (D) and der(1)t(1;14) (E). (F,G) Sanger sequencing shows the translocation junction of der(11)t(2;11) (F) and der(17)t(12;17) (G). Deletions (dashes), templated insertions (blue box), and microhomology regions (highlighted and underlined) are indicated.

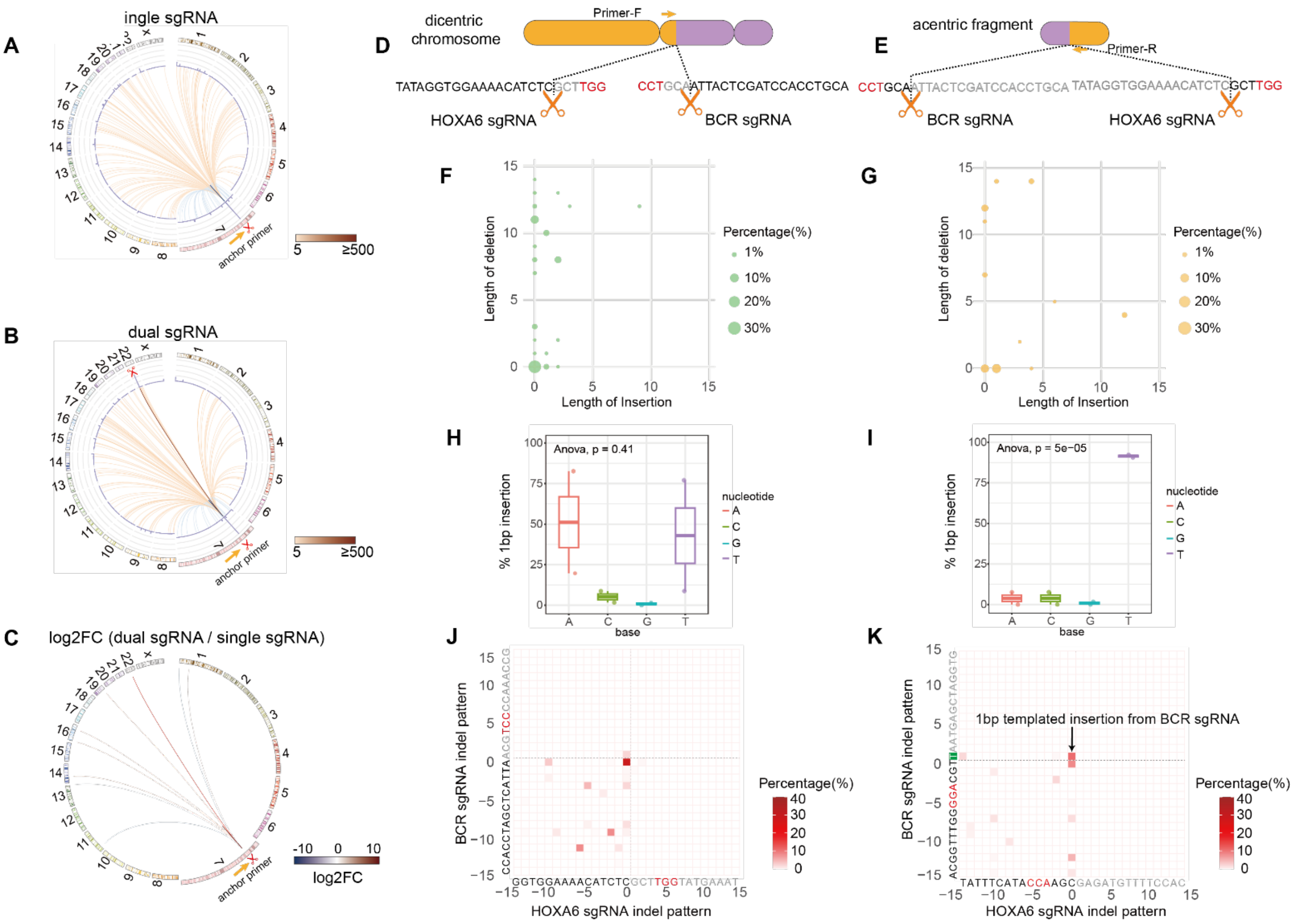


**Fig S2. Translocation landscapes and junctional features of unbalanced rearrangements.** (A-C) Genome-wide translocation distributions comparing single-sgRNA versus dual-sgRNA induction. Circos plots display inter-chromosomal translocations. Histograms represent translocation density across 500 bp genomic bins. Frequencies are normalized as reads per million (RPM). $Log_2$-transformed ratios of dual-sgRNA versus single-sgRNA highlight a pronounced increase in chromosomal translocations between specific loci. (D, E) Schematics of dicentric chromosome and acentric fragment formation, showing Cas9 cleavage sites, junction configurations, and anchor primer locations. (F, G) Bubble plot showing frequency and length distributions of deletions and insertions at unbalanced translocation junctions. Circle area reflects relative frequency. (H, I) Nucleotide composition of 1-bp insertions at dicentric and acentric junctions. Box plots show median and interquartile range (Q1-Q3), with the median indicated by a central line. (J, K) Heatmaps of length and frequency distributions of deletions and templated insertions (TINs) mapped to the *HOXA6* and *BCR* cleavage sites at dicentric and acentric junctions. Statistical significance in H, I is estimated by two-tailed one-way ANOVA.

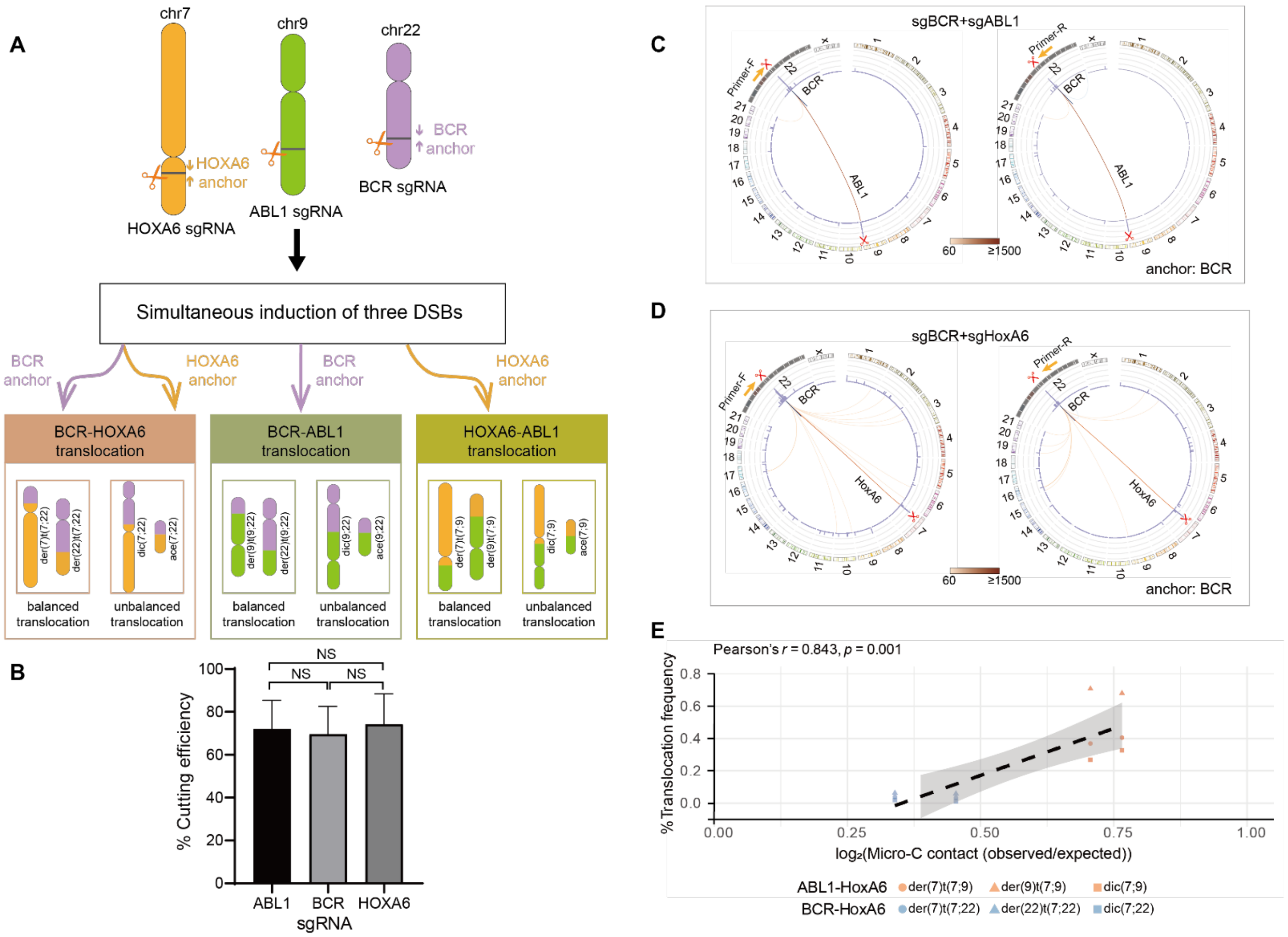


**Fig S3. Translocation mapping and spatial correlations programed by dual or triple sgRNAs.** (A) Schematic of chromosomal translocation outcomes and detection strategy following simultaneous induction of three DSBs at *BCR*, *ABL1*, and *HOXA6*. (B) Cleavage efficiencies of sgRNAs targeting *ABL1*, *BCR*, and *HOXA6*. Data are mean ± SD; two-tailed Student's t-test; NS, not significant. (C, D) Circos plots of genome-wide translocations with dual sgRNAs. Translocation densities are mapped to 500 bp bins. (E) Pearson correlation and linear regression (dotted line) between translocation frequencies and $\log_2$-transformed observed/expected Micro-C contact frequencies.

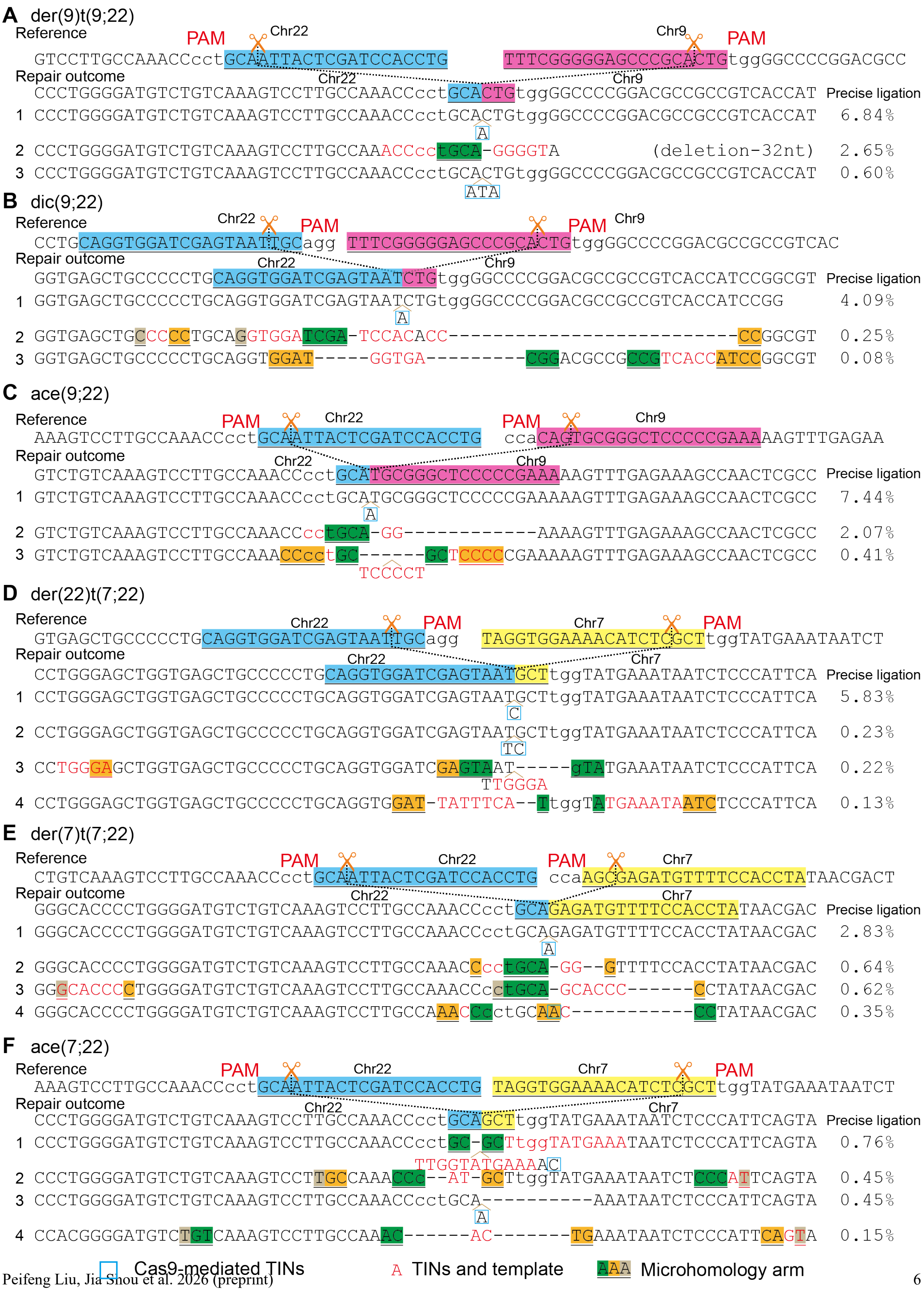
A der(9)t(9;22)
Reference
PAM Chr22 Chr9 PAM
GTCCTTGCCAAACCcctGCAATTACTCGATCCACCTG TTTCGGGGGAGCCCGCACTGtggGGCCCCGGACGCC
Repair outcome
Chr22 Chr9
CCCTGGGGATGTCTGTCAAAGTCCTTGCCAAACCcctGCACTGtggGGCCCCGGACGCCGCCGTCACCAT Precise ligation
1 CCCTGGGGATGTCTGTCAAAGTCCTTGCCAAACCcctGCACTGtggGGCCCCGGACGCCGCCGTCACCAT 6.84%
A
2 CCCTGGGGATGTCTGTCAAAGTCCTTGCCAAACCcctGCA-GGGGTA (deletion-32nt) 2.65%
3 CCCTGGGGATGTCTGTCAAAGTCCTTGCCAAACCcctGCACTGtggGGCCCCGGACGCCGCCGTCACCAT 0.60%
ATA
B dic(9;22)
Reference
Chr22 PAM PAM Chr9
CCTGCAGGTGGATCGAGTAATTGCagg TTTCGGGGGAGCCCGCACTGtggGGCCCCGGACGCCGCCGTCAC
Repair outcome
Chr22 Chr9
GGTGAGCTGCCCCCTGCAGGTGGATCGAGTAATCTGtggGGCCCCGGACGCCGCCGTCACCATCCGGCGT Precise ligation
1 GGTGAGCTGCCCCCTGCAGGTGGATCGAGTAATCTGtggGGCCCCGGACGCCGCCGTCACCATCCGG 4.09%
A
2 GGTGAGCTGCCCCTGCAGGTGGATCGA-TCCACACC---------------------------CCGGCGT 0.25%
3 GGTGAGCTGCCCCCTGCAGGTGGAT-----GGTGA---------CGGACGCCGCCGTCACCATCCGGCGT 0.08%
C ace(9;22)
Reference
PAM Chr22 PAM Chr9
AAAGTCCTTGCCAAACCcctGCAATTACTCGATCCACCTG ccaCAGTGCGGGCTCCCCCGAAAAAGTTTGAGAA
Repair outcome
Chr22 Chr9
GTCTGTCAAAGTCCTTGCCAAACCcctGCATGCGGGCTCCCCCGAAAAAGTTTGAGAAAGCCAACTCGCC Precise ligation
1 GTCTGTCAAAGTCCTTGCCAAACCcctGCATGCGGGCTCCCCCGAAAAAGTTTGAGAAAGCCAACTCGCC 7.44%
A
2 GTCTGTCAAAGTCCTTGCCAAACCcctGCA-GG------------AAAAGTTTGAGAAAGCCAACTCGCC 2.07%
3 GTCTGTCAAAGTCCTTGCCAAACCcctGC------GCTCCCCCGAAAAAGTTTGAGAAAGCCAACTCGCC 0.41%
TCCCCT
D der(22)t(7;22)
Reference
Chr22 PAM Chr7 PAM
GTGAGCTGCCCCCTGCAGGTGGATCGAGTAATTGCagg TAGGTGGAAAACATCTCGCTtggTATGAAATAATCT
Repair outcome
Chr22 Chr7
CCTGGGAGCTGGTGAGCTGCCCCCTGCAGGTGGATCGAGTAATGCTtggTATGAAATAATCTCCCATTCA Precise ligation
1 CCTGGGAGCTGGTGAGCTGCCCCCTGCAGGTGGATCGAGTAATGCTtggTATGAAATAATCTCCCATTCA 5.83%
C
2 CCTGGGAGCTGGTGAGCTGCCCCCTGCAGGTGGATCGAGTAATGCTtggTATGAAATAATCTCCCATTCA 0.23%
TC
3 CCTGGGAGCTGGTGAGCTGCCCCCTGCAGGTGGATCGAGTAAT-----gTATGAAATAATCTCCCATTCA 0.22%
TTGGGA
4 CCTGGGAGCTGGTGAGCTGCCCCCTGCAGGTGGAT-TATTTCA--TtggTATGAAATAATCTCCCATTCA 0.13%
E der(7)t(7;22)
Reference
PAM Chr22 PAM Chr7
CTGTCAAAGTCCTTGCCAAACCcctGCAATTACTCGATCCACCTG ccaAGCGAGATGTTTTCCACCTATAACGACT
Repair outcome
Chr22 Chr7
GGGCACCCCTGGGGATGTCTGTCAAAGTCCTTGCCAAACCcctGCAGAGATGTTTTCCACCTATAACGAC Precise ligation
1 GGGCACCCCTGGGGATGTCTGTCAAAGTCCTTGCCAAACCcctGCAGAGATGTTTTCCACCTATAACGAC 2.83%
A
2 GGGCACCCCTGGGGATGTCTGTCAAAGTCCTTGCCAAACCcctGCA-GG--GTTTTCCACCTATAACGAC 0.64%
3 GGGCACCCCTGGGGATGTCTGTCAAAGTCCTTGCCAAACCcctGCA-GCACCC------CCTATAACGAC 0.62%
4 GGGCACCCCTGGGGATGTCTGTCAAAGTCCTTGCCAAACCctGCAAC-----------CCTATAACGAC 0.35%
F ace(7;22)
Reference
PAM Chr22 Chr7 PAM
AAAGTCCTTGCCAAACCcctGCAATTACTCGATCCACCTG TAGGTGGAAAACATCTCGCTtggTATGAAATAATCT
Repair outcome
Chr22 Chr7
CCCTGGGGATGTCTGTCAAAGTCCTTGCCAAACCcctGCAGCTtggTATGAAATAATCTCCCATTCAGTA Precise ligation
1 CCCTGGGGATGTCTGTCAAAGTCCTTGCCAAACCcctGC-GCTtggTATGAAATAATCTCCCATTCAGTA 0.76%
TTGGTATGAAAAC
2 CCCTGGGGATGTCTGTCAAAGTCCTTGCCAAACCC--AT-GCTtggTATGAAATAATCTCCCATTCAGTA 0.45%
3 CCCTGGGGATGTCTGTCAAAGTCCTTGCCAAACCcctGCA----------AAATAATCTCCCATTCAGTA 0.45%
A
4 CCACGGGGATGTCTGTCAAAGTCCTTGCCAAAC------AC-------TGAAATAATCTCCCATTCAGTA 0.15%
Cas9-mediated TINs
A TINs and template
AAA Microhomology arm

**Fig S4. Representative junction features of Cas9-induced and Polθ-mediated templated insertions at translocation junctions.** (A-F) Representative alignments of translocation junctional sequences containing Cas9-induced TINs (Cas9-TINs) and Polθ-mediated TINs (Polθ-TINs) compared to precise ligation products. Translocation include der(9)t(9;22) (A), dic(9;22) (B), ace(9;22) (C), der(22)t(7;22) (D), der(7)t(7;22) (E), and ace(7;22) (F). Deletions (dashes), TINs (red), and microhomologies (highlighted and underlined) are indicated. Panels E and F show junctions containing co-occurring Cas9-TIN and Polθ-TIN within single sequence reads at translocation junctions.

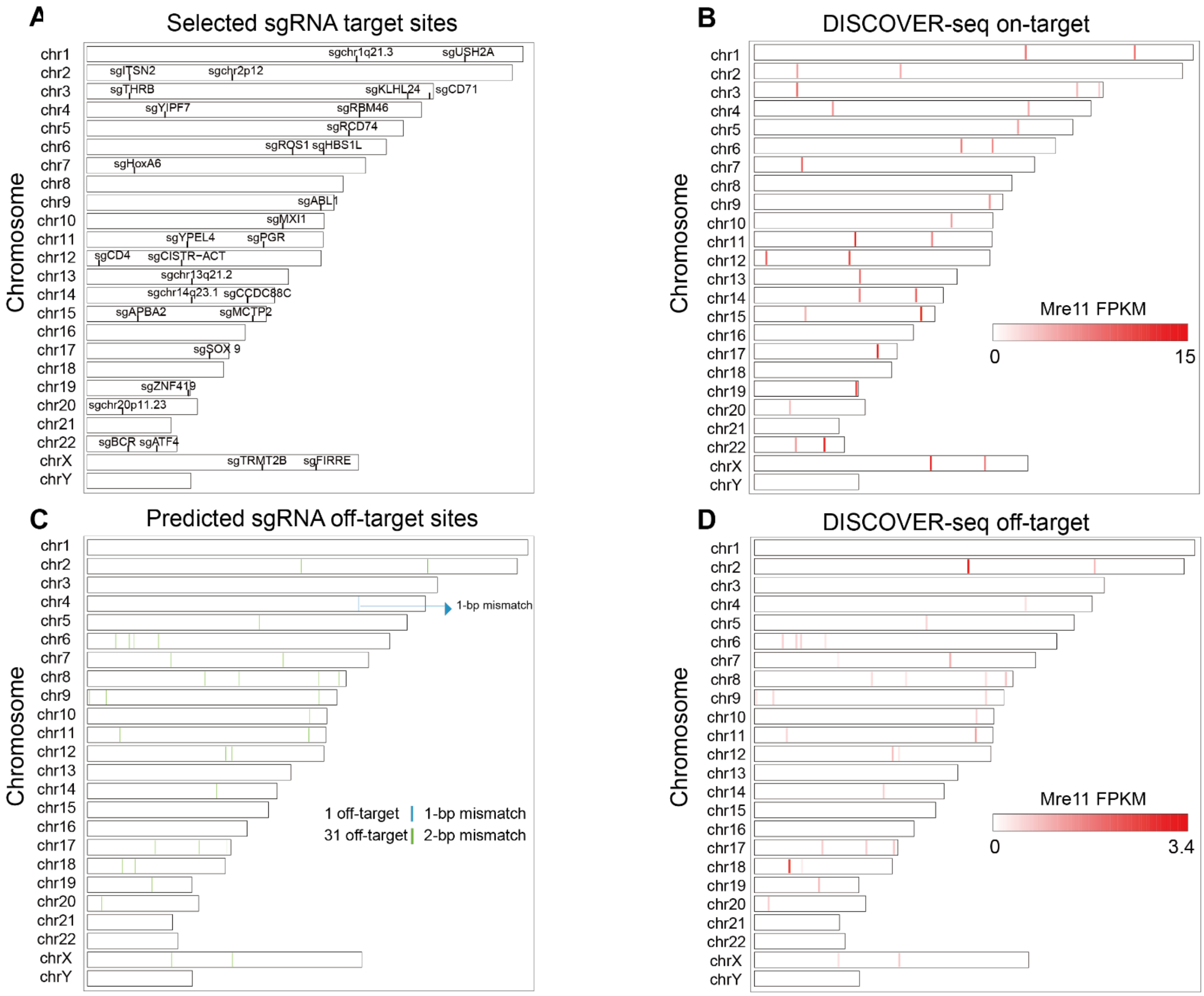


**Fig S5. DISCOVER-seq mapping of Cas9 cleavages at on-target and off-target sites.** (A) Genomic distribution of 31 on-target sites for DISCOVER-seq profiling. (B) Heatmap of MRE11 enrichments at the 31 on-target sites. Color intensity reflects the enrichment level. (C) Predicted off-target sites for the 31 sgRNAs by Cas-OFFinder. Sites with one mismatch are highlighted in blue, and sites with two mismatches are shown in green. (D) Heatmap of MRE11 enrichments at 32 off-target sites.

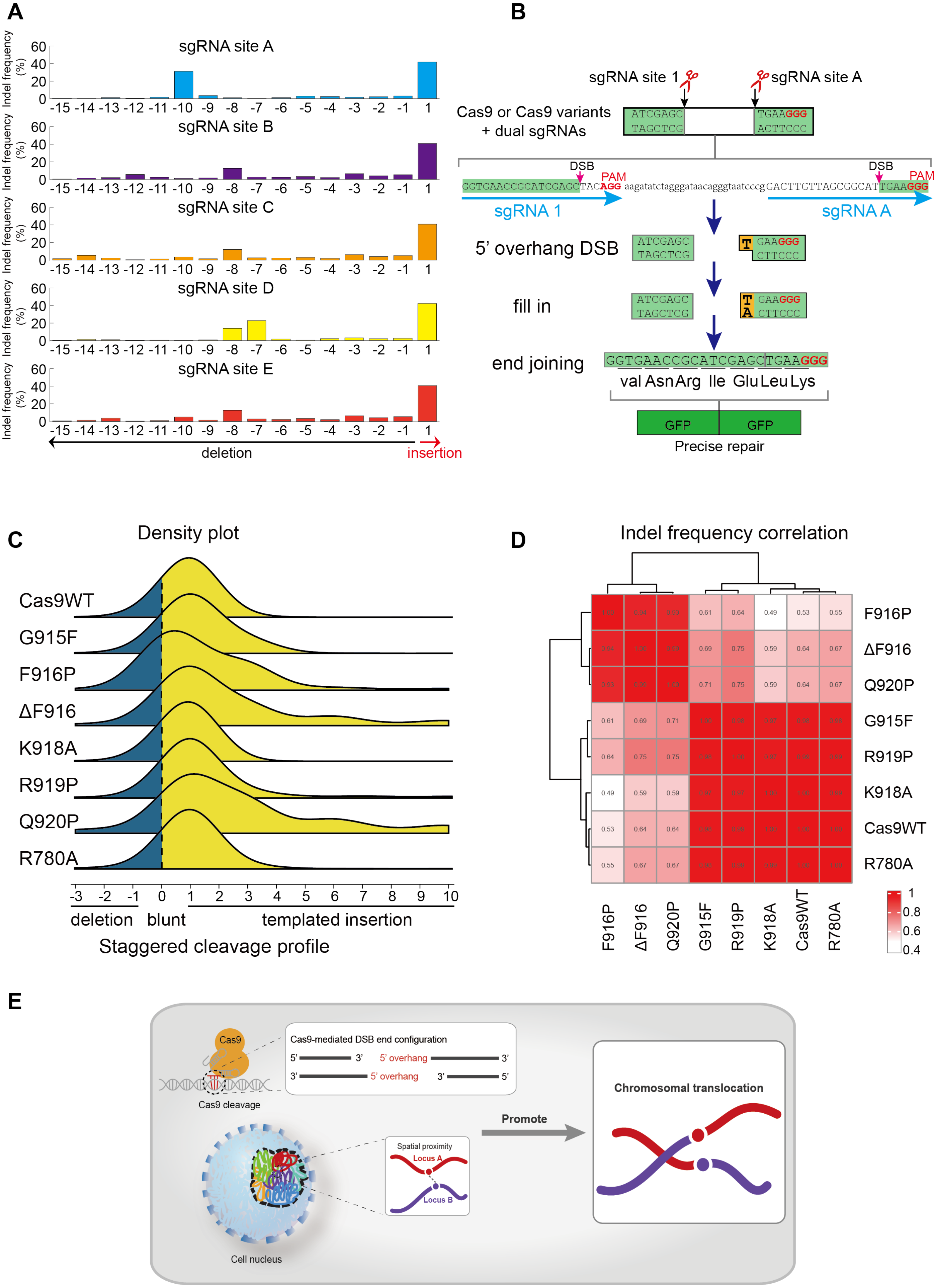

A
B
C
D
E
sgRNA site A
sgRNA site B
sgRNA site C
sgRNA site D
sgRNA site E
Indel frequency (%)
60
40
20
0
-15 -14 -13 -12 -11 -10 -9 -8 -7 -6 -5 -4 -3 -2 -1 1
deletion
insertion
sgRNA site 1
sgRNA site A
Cas9 or Cas9 variants + dual sgRNAs
ATCGAGC
TAGCTCG
TGAAGGG
ACTTCCC
DSB
PAM
GGTGAACCGCATCGAGCTACAGG aagatatctagggataacagggtaatcccg GACTTGTTAGCGGCATTGAAGGG
sgRNA 1
sgRNA A
5' overhang DSB
fill in
end joining
GGTGAACCGCATCGAGCTGAAGGG
val Asn Arg Ile Glu Leu Lys
GFP
GFP
Precise repair
Density plot
Cas9WT
G915F
F916P
ΔF916
K918A
R919P
Q920P
R780A
-3 -2 -1 0 1 2 3 4 5 6 7 8 9 10
deletion
blunt
templated insertion
Staggered cleavage profile
Indel frequency correlation
F916P
ΔF916
Q920P
G915F
R919P
K918A
Cas9WT
R780A
0.61 0.64 0.49 0.53 0.55
0.69 0.75 0.59 0.64 0.67
0.71 0.75 0.59 0.64 0.67
1
0.8
0.6
0.4
Cas9
Cas9 cleavage
Cas9-mediated DSB end configuration
5' 3' 5' overhang 3'
3' 5' overhang 3' 5'
Cell nucleus
Spatial proximity
Locus A
Locus B
Promote
Chromosomal translocation

**Fig S6. Engineered Cas9 variants modulate end geometry and repair outcomes.** (A) InDelphi-predicted indel profiles for five candidate sgRNAs evaluated for reporter design; sgRNA site A was selected for subsequent experiments. (B) Schematic of the dual-sgRNA GFP reporter system. GFP restoration requires a precise +1 bp templated insertion generated from Cas9-mediated staggered cleavage. (C) Density plot of junctional indel distributions by deep sequencing. The black vertical line marks the blunt-end cleavage position. Positive values indicate insertions and negative values represent deletions. Note that repair outcomes of F916P, ΔF916 and Q920P are shifted toward longer templated insertions. (D) Hierarchical clustering of junctional indel patterns across Cas9WT and engineered variants based on Pearson correlation coefficients. (E) Proposed model in which 3D genome architecture constrains DSB encounter probability while the configuration of Cas9-induced DSB end governs repair compatibility, jointly determining the frequency and precision of chromosomal translocations.
